# Cortical excitation-inhibition ratio mediates the effect of pre-attentive auditory processing deficits on interpersonal difficulties

**DOI:** 10.1101/728204

**Authors:** Talitha C. Ford, Will Woods, Peter G. Enticott, David P. Crewther

## Abstract

Several lines of evidence identify aberrant excitatory-inhibitory neural processes across autism and schizophrenia spectrum disorders, particularly within the psychosocial domain. Such neural processes include increased excitatory glutamate and reduced inhibitory GABA concentrations, which may affect auditory pre-attentive processing as indexed by the mismatch negativity (MMN); thus, an excitation-inhibition imbalance might lead to aberrant MMN, which might in turn drive the relationship between the MMN and psychosocial difficulties. This research has the potential to enhance the neurochemical understanding of the relationship between electrophysiology (MMN) and behavioural/clinical measures (psychosocial difficulties).

Thirty-eight adults (18 male, 18-40 years) completed the Schizotypal Personality Questionnaire (SPQ) and Autism-Spectrum Quotient (AQ). Glutamate and GABA concentrations in bilateral superior temporal cortex (STC) were quantified using proton magnetic resonance spectroscopy (1H-MRS) while auditory MMN to a duration deviant was measured with magnetoencephalography. Spearman correlations probed the relationships between STC glutamate/GABA ratios, MMN amplitude and latency, and AQ and SPQ dimensions. Mediation effects of glutamate/GABA ratios on the relationship between MMN and AQ-SPQ dimensions were probed using causal mediation analysis.

Only SPQ-interpersonal and AQ-communication were significantly correlated with right hemisphere glutamate/GABA ratios and MMN latency (*p*s<.05), which were themselves correlated (*p*=.038). Two mediation models were investigated, with right MMN latency as predictor and SPQ-interpersonal and AQ-communication as outcome variables. Right STC glutamate/GABA ratios significantly mediated the relationship between MMN latency and SPQ-interpersonal scores (ß=86.6, *p*=.033), but only partially mediated the relationship between MMN latency and AQ-communication scores (ß=21.0, *p*=.093).

These findings support the growing body of literature pointing toward an excitation-inhibition imbalance that is central to psychosocial functioning across multi-dimensional spectrum disorders, such as autism and schizophrenia, and provides neurochemical indicators of the processes that underlie psychosocial dysfunction.

---

The diversity of risk-genes, neurobiological abnormalities, and environmental factors that contribute to the symptomatology of autism and schizophrenia lead to its multidimensionality and heterogeneity (Foss-Feig et al., 2017). Several lines of evidence point toward overlapping symptomatology between autism and schizophrenia, particularly in relation to social and communication dysfunction (Canitano and Pallagrosi, 2017; Chisholm et al., 2015; Chung et al., 2014; Ford and Crewther, 2014; Shi et al., 2017), and converging neurobiological and genetic evidence further suggests that overlapping symptoms have similar underlying mechanisms (Canitano and Pallagrosi, 2017; Chisholm et al., 2015; Foss-Feig et al., 2017; Pinkham et al., 2008; Sugranyes et al., 2011). The mismatch negativity (MMN), a pre-attentive auditory processing mechanism, has been implicated across both conditions (Bridwell et al., 2014; Chien et al., 2018; Kargel et al., 2014; Schwartz et al., 2018), particularly in relation to poor psychosocial functioning (Canitano and Pallagrosi, 2017; Chien et al., 2018; Fan and Cheng, 2014; Greenwood et al., 2018; Kargel et al., 2014; Lee et al., 2014; Wynn et al., 2010).

The auditory MMN is elicited in the event of a deviation to the constant auditory environment (Naatanen et al., 2012), and is thought to be governed by excitatory glutamatergic neurotransmitter signalling through NMDA receptor channels in auditory and frontal brain regions (Javitt et al., 1996; Rosburg and Kreitschmann-Andermahr, 2016; Rowland et al., 2016; Wacongne et al., 2012). In fact, disruption to normal NMDA receptor functioning through pharmaceutical agents acting upon glutamatergic or GABAergic processes (e.g., ketamine, lorazepam) leads to a reduced in MMN amplitude and a delayed MMN latency (Kreitschmann-Andermahr et al., 2001; Rosburg and Kreitschmann-Andermahr, 2016; Umbricht et al., 2002). Acute glycine administration, which increases NMDA receptor-mediated glutamate function, has been shown to normalise MMN amplitude and latency in schizophrenia patients, in combination with improved negative/psychosocial symptom outcomes (Greenwood et al., 2018). Inhibitory GABA functionality has also been implicated in the MMN, with the GABA agonist lorazepam shown to reduce MMN amplitude and delay MMN latency in a small sample of healthy young adults (Rosburg et al., 2004).

Ketamine, an NMDA antagonist, has also been shown to increase glutamate and glutamine (a putative marker of glutamate transmitter release) concentrations in the anterior cingulate cortex (Rowland et al., 2016; Stone et al., 2012), and a recent meta-analysis demonstrated increased glutamate and glutamate+glutamine (Glx) concentrations for schizophrenia patients across the basal ganglia, and temporal and frontal regions (Merritt et al., 2016).

NMDA receptor dysfunction, as well as dysregulation of excitatory glutamatergic and inhibitory GABA neurotransmission, has been implicated across both autism and schizophrenia spectrum disorders (Ajram et al., 2019; Canitano and Pallagrosi, 2017; Cellot and Cherubini, 2014; Deidda et al., 2014; Fatemi, 2008; Fatemi et al., 2009; Foss-Feig et al., 2017; Greenwood et al., 2018; Merritt et al., 2016; Moghaddam and Javitt, 2012; Rojas, 2014), and poor psychosocial functioning (Cochran et al., 2015; Ford et al., 2017a; Greenwood et al., 2018; Han et al., 2014; Hegarty et al., 2018; Thiebes et al., 2017; Yizhar et al., 2011). However, the extent to which the relationship between the MMN and psychosocial deficits is modulated by excitatory-inhibitory processes is not yet known.

In healthy populations, higher Glx concentrations in the superior temporal cortex have been associated with shorter MMN latency and slightly increased MMN amplitude (Kompus et al., 2015), while hippocampal glutamate/creatine ratios were associated with MMN amplitudes measured from temporal (not frontal or central) electrode sites (Chitty et al., 2015), and frontally recorded MMN amplitude was not associated with anterior cingulate glutamate or GABA concentrations (Rowland et al., 2016). These relationships were more specific to deviations in stimulus duration, not frequency, which is thought to reflect the complexity of duration tone changes, compared to frequency (Todd et al., 2008, 2003). Furthermore, deviations in the duration of tones might rely on multiple system integration, or a specific neural system, that is governed by glutamatergic neurotransmission (Kompus et al., 2015). Duration tone deviants may, therefore, rely more heavily on multisystem integration, and be more sensitive in detecting differences in psychiatric disorders (Kompus et al., 2015; Todd et al., 2003).

In psychiatric populations, reduced glutamate and GABA concentrations in the anterior cingulate have been associated with reduced frontally recorded duration deviant MMN for schizophrenia patients (Rowland et al., 2016), and hippocampal glutamate/creatine ratios were associated with duration MMN amplitudes from temporal (not frontal or central) electrode sites in healthy controls, but not bipolar patients (Chitty et al., 2015). A more recent combined schizophrenia and bipolar study reported that increased frontal and right temporal duration MMN was associated with higher anterior cingulate glutamate/creatine ratios for schizophrenia patients only, while reduced MMN peak amplitude at frontal and temporal sites was associated with increased hippocampal glutamate/creatine, albeit not significantly (Kaur et al., 2019). It is important to note that these studies did not report on glutamate and GABA concentrations in relation to MMN latency. Clearly the relationship between MMN amplitudes and glutamate and GABA concentrations is highly variable across brains regions, and may be affected by sample characteristics, such as medication use, symptom severity and age; thus, investigating the interrelationship of MMN and glutamate/GABA ratios on specific symptom traits within a unmedicated population, while targeting a socially relevant brain region, may provide key insights into the role of excitation-inhibition in autism and schizophrenia symptomatology.

To that end, we investigated the relationship between the auditory cortex generated MMN measured with magnetoencephalography (MEG), the mismatch field (MMF), and the core subclinical autism and schizophrenia spectrum domains, further probing the extent to which superior temporal glutamate/GABA ratios (as measured via proton magnetic resonance spectroscopy [1H-MRS]) mediate these relationships. Given evidence based on NMDA receptor dysfunction (Kreitschmann-Andermahr et al., 2001; Rosburg and Kreitschmann-Andermahr, 2016; Stone et al., 2012; Umbricht et al., 2002), it was hypothesised that higher glutamate/GABA ratios would be associated with delayed MMF latencies and smaller MMF amplitudes. Based on previous research by our group (Ford et al., 2017b, 2017c), we hypothesised that delayed MMF latency would be associated with psychosocial symptom domains, particularly over the right hemisphere. We also predicted that a higher ratio of glutamate/GABA in the right superior temporal cortex (STC) would be associated with poorer psychosocial functioning. Finally, we expected to elucidate a mediating effect of glutamate/GABA ratios on any relationships between MMN latency and psychosocial subscale scores.

## Materials and Methods

Ethical approval for this study was granted by the Swinburne University Human Research Ethics Committee (2011/033 Series C(d)) in accordance with the Declaration of Helsinki. All participants provided written informed consent prior to commencing the study.

### Participants

Participant recruitment and data collection procedures for this study have been described previously (Ford et al., 2017a, 2017d; Ford and Crewther, 2014). Initially, 1,678 adults aged 18 to 40 years (428 male, 1250 female) were recruited to complete an online questionnaire consisting of the autism-spectrum quotient (AQ) and the schizotypal personality questionnaire (SPQ). The AQ and SPQ subscales were entered into an exploratory factor analysis to identify latent factors that exist within the autism-schizotypy spectra; a combined autism-schizotypy factor was revealed and termed Social Disorganisation (Ford and Crewther, 2014). Participants scoring in the top and bottom 20% of Social Disorganisation scores were invited to participate in a follow-up neuroimaging study involving MEG and 1H-MRS. The first 10 male and 10 female responders from each end of the Social Disorganisation distribution were recruited. In total, 38 participants took part in the neuroimaging study (20 female, 18 male).

One female participant was excluded due to reporting current psychoactive medications; of the remaining participants, five self-reported a previous psychiatric condition (3 depression, 1 bipolar, 1 anorexia). No participant reported a previous or current autism or schizophrenia spectrum condition, and none were affected by a psychiatric condition at the time of the study. All participants reported being free of illicit drug and cigarette effects at the time of the MEG and MRI scans.

Given the ovulation phase of the female menstrual cycle has been shown to affect GABA concentrations (De Bondt et al., 2015), the number of days post-menses onset was obtained. There was no relationship between and menstrual day and left or right STC GABA levels (*p* > .1; see the supplement of Ford et al., 2019 for further details).

### Materials

The 50-item AQ was used to assess autistic tendency. Items are collapsed into five subscales: social skill, communication, attention switching, attention to detail and imagination (Baron-Cohen et al., 2001). The 74-item SPQ was used to measure schizotypal personality traits (Raine, 1991). Three superordinate dimensions of schizophrenia spectrum disorder encapsulate nine SPQ subscales: ideas of reference, odd beliefs, unusual perceptual experiences, and suspiciousness (cognitive-perceptual features); social anxiety, no close friends and constricted affect (interpersonal features); odd behaviour and odd speech (disorganised features). AQ and SPQ items were pseudo-randomised and presented on a 4-point Likert scale from 1 (*strongly disagree*) to 4 (*strongly agree*) in order to improve reliability (Ford and Crewther, 2014; Wuthrich and Bates, 2005).

General intelligence was obtained using the 12-item short version of the Ravens Advanced Progressive Matrices (RAPM; Arthur and Day, 1994; Raven et al., 1998). Particiants completed as many of the 12 matrices as possible within 10 minutes, with the number of complete and correct responses recorded as participants total RAPM score.

### Neuroimaging procedures

#### ^1^H-MRS data acquisition and processing

The ^1^H-MRS protocol has been detailed previously (Ford et al., 2017a, 2017b). Briefly, T1-weighted images were recorded from a 3T Siemens TIM Trio whole-body magnetic resonance imaging system (Erlangen, Germany) equipped with a 32-channel head coil. T1-weighted images were acquired using an MPRage pulse sequence with an inversion recovery (176 slices, slice thickness = 1.0mm, voxel resolution = 1.0mm^3^, TR = 1900ms, TE = 2.52ms, TI = 900ms, bandwidth = 170Hz/Px, flip angle = 9°, field of view 350mm x 263mm x 350mm, orientation sagittal, acquisition time = ∼5min).

The T1 image was used to position the left and the right STC voxels (20×30×20mm). STC voxels were placed along the posterior portion of the superior temporal gyrus, encapsulating the primary auditory cortex, temporoparietal junction and inferior portion of the inferior parietal lobe; see Figure 1 for example voxel placement and Supplementary Figure 1 for mean voxel placement on a standard MNI structural image. Voxels A PRESS sequence with an echo time of 80ms was employed to quantify glutamate concentrations from temporal voxels (Schubert et al., 2004)(TE = 80ms, TR = 2000ms, bandwidth = 1200Hz, 80 averages, 8 spectral water averages, acquisition time = 2 min 48 sec). A MEGA-PRESS (Mescher et al., 1998) editing sequence was applied to quantify temporal GABA concentrations (TE = 68ms, TR = 1500ms, bandwidth = 1000Hz, edit-on pulse frequency = 1.9ppm, edit-off pulse frequency = 7.5ppm, edit pulse bandwidth = 44Hz, 120 averages, 12 spectral unsuppressed water averages, duration = 6min 6 sec). As an edit-off pulse at 7.5ppm does not suppress macromolecular concentrations at 3.0ppm, the spectra included macromolecules (GABA+). For all scans, chemical shift selective (CHESS; Haas, 1986) water suppression was applied, and automatic and manual shimming was conducted to achieve full-width half maximum (FWHM) of < 20. Right and left temporal voxel PRESS and MEGA-PRESS spectra averages are illustrated in Figure 1.

**Figure 1.**
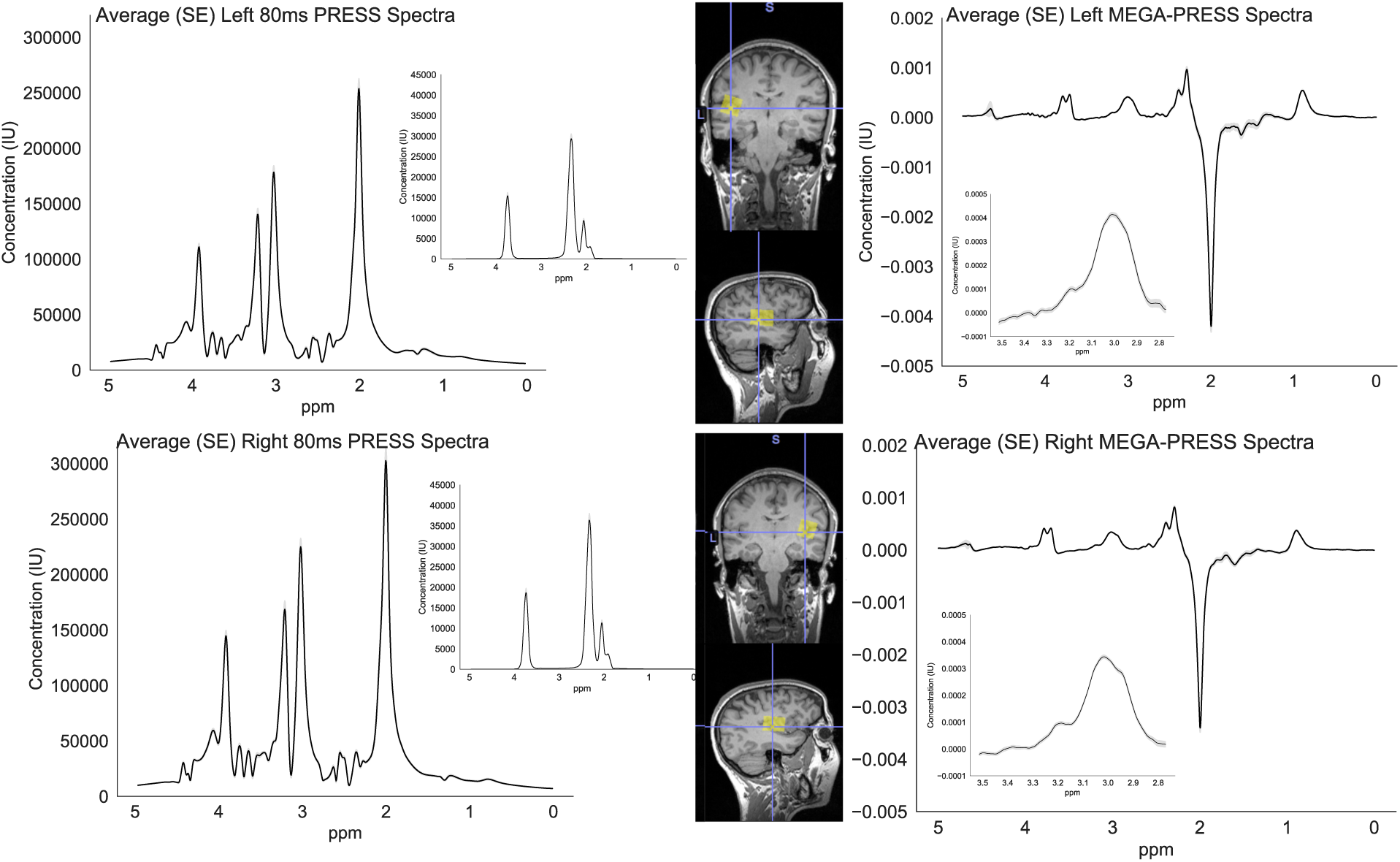
Average PRESS and MEGA-PRESS spectra and voxel placement. Left panel: average spectra for PRESS sequence for left (top) and right (bottom) superior temporal voxels, with standard error shown in grey shading. Middle panel: the isolated glutamate peak is presented in the inset. Voxel placement for left (top) and right (bottom) 20×30×20mm superior temporal voxels. Right panel: average spectra for MEGA-PRESS sequence for left (top) and right (bottom) superior temporal voxels, with standard error shown in grey shading. The isolated GABA+ peak is presented in the inset.

PRESS and MEGA-PRESS data were processed using Tarquin (version 4.3.7; Wilson et al., 2011) and Gannet’s GABA analysis toolkit for Matlab (version 2.0; Edden et al., 2014), respectively. For quantification of glutamate with Tarquin, a non-negative least-squares projection of a parametrised basis set was used to estimate signal amplitude in the time-domain, and eddy current correction was applied. The unsuppressed water spectra were used as a reference for metabolite quantification, resulting in glutamate concentrations in institutional units (IU). The data of one participant was excluded due poor data quality (left water frequency standard deviation = −6.88, right FWHM = 14.22), and no right hemisphere data was recorded for one participant. All remaining participants’ data had adequate fit (*n* = 35; see Supplementary Table 1>).

**Table 1.** Mean and standard error (SE) for demographic, MEG and 1H-MRS measures.

|  | Mean (SE) |  |  |  |
| --- | --- | --- | --- | --- |
|  | Female (*n*=17) | Male (*n*=16) |  |  |
| Age in years | 22.41 (1.13) | 23.5 (1.43) |  |  |
| RAPM | 7.41 (.54) | 6.5 (.60) |  |  |
| AQ Social Skills | 3.94 (.76) | 3.75 (.60) |  |  |
| AQ Communication | 3.77 (.53) | 3.62 (.67) |  |  |
| AQ Attention Switching | 3.71 (.67) | 3.88 (.75) |  |  |
| AQ Attention to detail | 5.06 (.30) | 5.5 (.47) |  |  |
| AQ Imagination | 3.23 (.33) | 3.69 (.45) |  |  |
| AQ Total | 115.4 (5.40) | 111.9 (5.99) |  |  |
| SPQ Cognitive Perceptual features | 8.59 (1.45) | 9.44 (1.29) |  |  |
| SPQ Interpersonal features | 9.59 (1.68) | 11.25 (1.78) |  |  |
| SPQ Disorganisation features | 5.29 (.89) | 7.69 (1.16)* |  |  |
| SPQ Total | 157.6 (10.92) | 147.8 (10.08) |  |  |
|  | Left hemisphere |  | Right hemisphere |  |
|  | Female (*n*=16) | Male (*n*=16) | Female (*n*=17) | Male (*n*=16) |
| MMF amplitude (fT) | 2.21 (.34) | 2.25 (.32) | 2.42 (.23) | 2.41 (.26) |
| MMF latency (sec) | .174 (.005) | .174 (.004) | .174 (.004) | .176 (.004) |
| GABA+ (IU) | 2.68 (.20) | 2.41 (.13) | 2.16 (.16) | 3.00 (.38) |
| Glutamate (IU) | 6.04 (.26) | 6.15 (.20) | 5.09 (.15) | 4.83 (.20) |
| Glx (IU) | 6.72 (0.32) | 7.03 (0.26) | 5.75 (0.22) | 5.51 (0.24) |
| Glutamate/GABA ratio | 2.48 (.25) | 2.66 (.17) | 2.54 (.18) | 2.03 (.25) |

Note: \* *p* < .05. RAPM = Raven's Advanced Progressive Matrices, AQ = autism spectrum quotient, SPQ = schizotypal personality
questionnaire, MMF = mismatch field, fT = femtotesla, sec = seconds, IU = institutional units, Glx = glutamate + glutamine.

For quantification of GABA with Gannet, edit-on and edit-off spectra were subtracted to reveal GABA+ concentration at 3.00ppm relative water, resulting in GABA+ concentrations in IU (Mullins et al., 2014). All GABA spectra had a fit error of < 15% and creatine line-width < 11Hz. An in-house voxel tissue segmentation Matlab script was used to quantify the amount of grey matter, white matter and cerebro-spinal fluid in each superior temporal voxel for each participant. Glutamate and GABA+ concentrations were then corrected for individual voxel tissue composition differences using the formula given by Harris et al. (2015). The ratio of glutamate/GABA+ was calculated as a proxy for cortical excitation-inhibition (Ford et al., 2017b; Rubenstein and Merzenich, 2003).

#### MEG data acquisition and processing

The MEG protocol utilised in this study has been reported in detail previously (Ford et al., 2017c). In brief, continuous MEG data were collected using a whole-head 306 channel Elekta Neuromag® TRIUX magnetometer system (Helsinki, Finland; Elekta, 2016) within a magnetically shielded room, with external active shielding on, at a rate of 1000Hz with a .1Hz high-pass filter. Five continuous head position 1 indicators tracked head position in relation to magnetometers, and head shape was digitised using a Polhemus FASTRACK system (Polhemus Inc., Colchester, VT, United States of America). Blink artefact and heart rate artefact were detected using electrodes placed above and below the right eye, and on the right wrist, respectively. A reference electrode was positioned on the right elbow. Participants were seated and watched a silent film while passive MMN stimuli were presented binaurally through Etymotic headphones. Stimuli consisted of three blocks of standard tones (50ms, 1000Hz, 85% of stimuli) randomly interrupted by duration deviants (100ms, 1000Hz, 15% of stimuli). The inter-stimulus interval was 500ms (offset to onset) and each block lasted 4–5 min. Stimuli were created using VPixx version 3.0 with DataPixx (VPixx Technologies Inc., 2016) and calibrated with a Jay Cam sound pressure level meter to 85 dB SPL.

For all raw MEG data, MaxFilter (version 2.1; Helsinki, Finland: Elekta, 2016) with default parameters was utilised for temporal signal space separation (tSSS) filtering with initial head position and bad channels removed. The data of one male participant were corrupted. Remaining bad channels, and highly variable stimulus epochs, were identified by eye. Original MEG data were then passed through MaxFilter for a second time excluding the additional bad channels.

Pre- and post-processing were conducted using MNE Python (Gramfort et al., 2014, 2013). Maxfiltered data were low-pass filtered (40Hz) and epoched (−200ms – 500ms), and epochs were baseline corrected with the pre-stimulus interval (−200ms – 0ms). Standard and deviant stimulus epochs were averaged for each participant (171 deviant stimuli per participant, on average), with only standard stimuli preceding a deviant included (176 standard stimuli per participant, on average), the difference calculated by subtracting the average standard waveform from the average deviant waveform. The first positive magnetic field power peak between 100ms and 200ms was identified as the MMF. The magnetometer with the strongest MMF (and largest signal to noise) in both the left and right temporal regions was identified for MMF peak power and latency analysis (right magnetometer: MEG2411, left magnetometer: MEG1621; Figure 2). Participant MMF data were excluded where MMF power did not exceed baseline.

**Figure 2.**
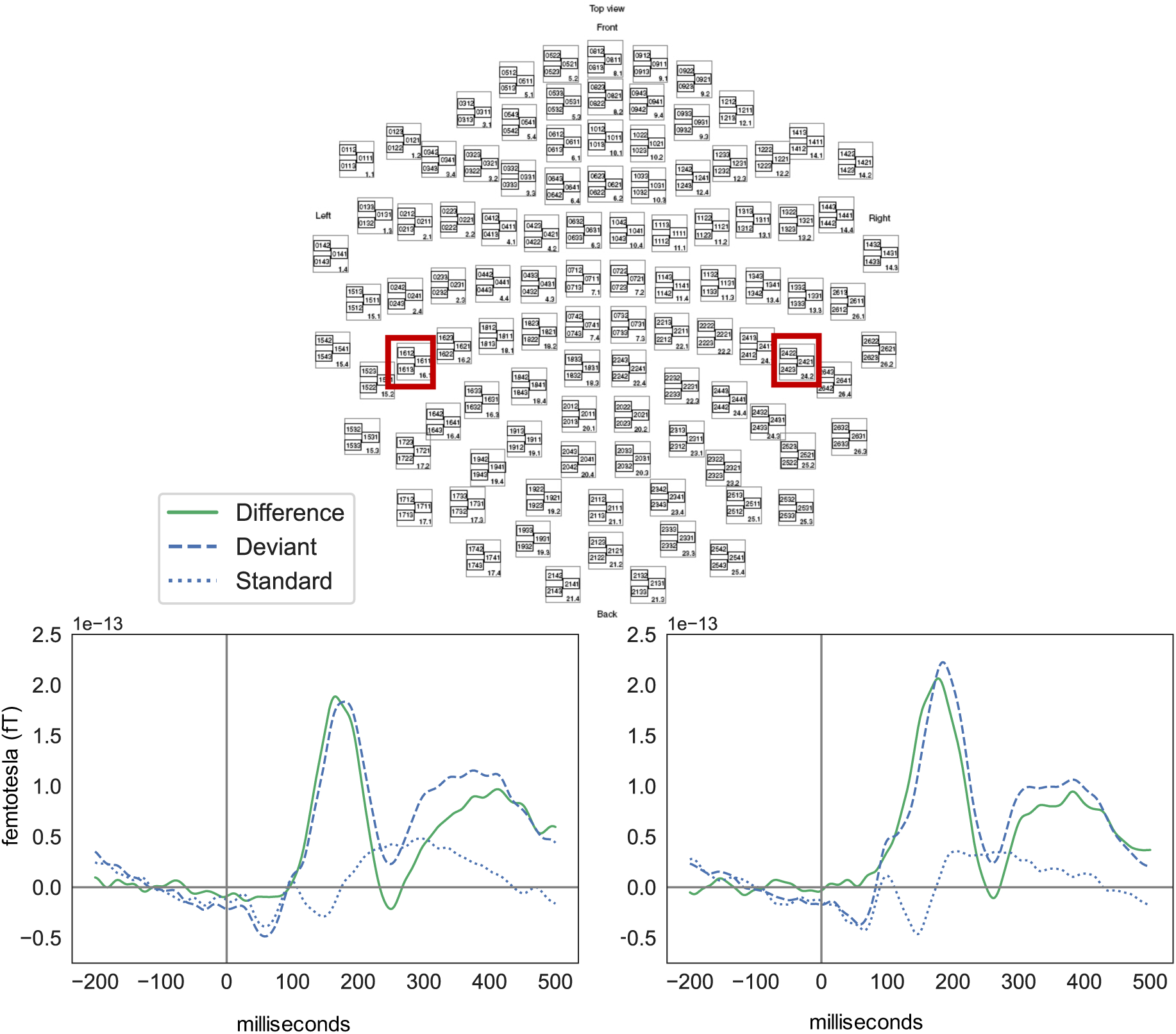
Average standard, deviant and difference waves across participants. Top: magnetoencephalography sensor array, with the red box indicating the peak left and right hemisphere sensors localised over the auditory cortex. Bottom left: Average left auditory response to standard and duration deviant tones, and the difference wave demonstrating a mismatch field (MMF) between 100-200ms. Bottom right: Average right auditory response to standard and duration deviant tones, and the difference wave demonstrating a MMF between 100-200ms.

### Statistical Analysis

Data were first screened for outliers, with STG GABA+ data excluded from two participants (left *n* = 1, right *n* = 1). The MMF was not detected on four occasions (left *n* = 2, right *n* = 2), and was thus removed from further statistical analyses. Statistical analyses were then conducted on the remaining datasets where 1H-MRS and MEG data were available for each hemisphere (left *n* = 32, right *n* = 33). All analyses were conducted using R (R Core Team, 2014), and the code is available at www.doi.org/10.13140/RG.2.2.29277.56800.

Spearman’s rank order (*ρ*) correlation coefficients were calculated to investigate relationships between the AQ and SPQ subscales, left and right MMF amplitude and latency, and left and right glutamate/GABA+ ratio, and to probe potential mediation models. Only correlation coefficients greater than *ρ* = .30 were considered (Tabachnick and Fidell, 2013). Spearman correlation coefficients for the relationships between the left temporal MMF and left glutamate/GABA ratio were below the *ρ* = .30 threshold (see Table 2), thus no left hemisphere mediation models were explored.

**Table 2.** Spearman’s rank order (*ρ*) correlations between AQ subscales and SPQ dimensions, and MEG and 1H-MRS measures

|  | MMF Latency |  | Glutamate/GABA+ |  |
| --- | --- | --- | --- | --- |
| | $\rho$ | $p$-value | $\rho$ | $p$-value |
| *Right hemisphere* |  |  |  |  |
| MMF amplitude | .151 | .403 | .162 | .367 |
| MMF latency |  |  | **.368\*** | **.035** |
| Social Skills | .283 | .111 | .476** | .005 |
| Communication | **.529\*\*** | **.002** | **.408\*** | **.019** |
| Attention Switching | .332 | .059 | .199 | .268 |
| Attention to detail | .356* | .042 | .191 | .288 |
| Imagination | .401* | .021 | .198 | .270 |
| Cognitive Perceptual features | .374* | .032 | .284 | .109 |
| Interpersonal features | **.391\*** | **.024** | **.453\*\*** | **.008** |
| Disorganised features | .537** | .001 | .237 | .184 |
| *Left hemisphere* |  |  |  |  |
| MMF amplitude | -.118 | .521 | -.001 | .994 |
| MMF latency |  |  | -.119 | .517 |
| Social Skills | .283 | .116 | -.087 | .635 |
| Communication | .165 | .367 | -.199 | .275 |
| Attention Switching | .352* | .048 | -.294 | .102 |
| Attention to detail | -.111 | .545 | .024 | .897 |
| Imagination | .156 | .392 | -.318 | .076 |
| Cognitive Perceptual features | .393* | .026 | -.119 | .518 |
| Interpersonal features | .279 | .123 | -.116 | .525 |
| Disorganised features | .175 | .338 | -.211 | .247 |

Notes: \* $p < .05$ , \*\* $p < .01$ , \*\*\* $p < .001$ , MMF = mismatch field, bold face indicates moderate and significant correlations across MMF latency and glutamate/GABA+ ratio.

Due to the moderate and significant correlations between right MMF latency, right glutamate/GABA+ ratio, AQ communication and SPQ interpersonal features (*ρ* > .30, *p* < .05), a mediation model was developed to investigate the extent to which right STC glutamate/GABA+ mediated the relationship between right MMF latency and AQ communication and SPQ interpersonal features. A series of regressions were used to fit models for the effect of the predictor on the mediator (*path a*), the mediator on the outcome (*path b*), and the predictor on the outcome variables (*path c*), as well as the effect of the predictor on the outcome controlling for the mediator (*path c’*). Age, sex and RAPM scores were entered as covariates. Linear models for *path a* and *path c’* were then utilised to test the significance of the mediation effect using Nonparametric Bootstrap Confidence Intervals with the Percentile Method from the *mediation* package in R (Tingley et al., 2014). Data were bootstrapped 10,000 times to calculate the average causal mediation effect (ACME); ACME was considered significant when 95% confidence intervals did not include zero.

## Results

Descriptive statistics for AQ subscales, SPQ dimensions, MMF amplitude and latency, and glutamate/GABA+ ratio are presented in Table 1. There were no sex differences in demographic, MMF or 1H-MRS variables (*p* > .1), except that SPQ Disorganised features were higher for males than females (*p* = .030).

In the right hemisphere, there was a significant moderate correlation between glutamate/GABA+ ratio and MMF latency, but not MMF amplitude. There were also significant correlations between MMF latency and AQ communication, AQ imagination, SPQ interpersonal features and SPQ disorganised features, and between glutamate/GABA+ ratio and AQ social skills, AQ communication and SPQ interpersonal features (see Table 2). Correlations between glutamate, Glx and GABA, and MMF amplitude and latency, are presented in Supplementary Table 2.

Given both MMF latency and glutamate/GABA+ ratio were significantly correlated with AQ communication and SPQ interpersonal features, separate mediation models were developed to investigate the extent to which glutamate/GABA+ ratio mediated the relationship between MMF latency and AQ communication, and MMF latency and SPQ interpersonal features. As illustrated in Figure 3, when controlling for age, sex and RAPM scores, MMF latency significantly predicted communication scores (ß= 74.12, *t* = 3.18[23.27], *p* = .004), and glutamate/GABA+ ratio (ß = 22.17, *t* = 2.43[9.14], *p* = .022). The relationship between MMF latency and communication was significant, albeit weaker when controlling for the effect of glutamate/GABA+ ratio, suggesting that glutamate/GABA+ ratio partially mediates this relationship (ß = 57.18, *t* = 2.30[24.87], *p* = .029). Nonparametric causal mediation analysis indicated that this mediating effect (ACME) was not significant (ß = 16.94, 95%CI[-7.49, 47.30], *p* = .161).

**Figure 3.**
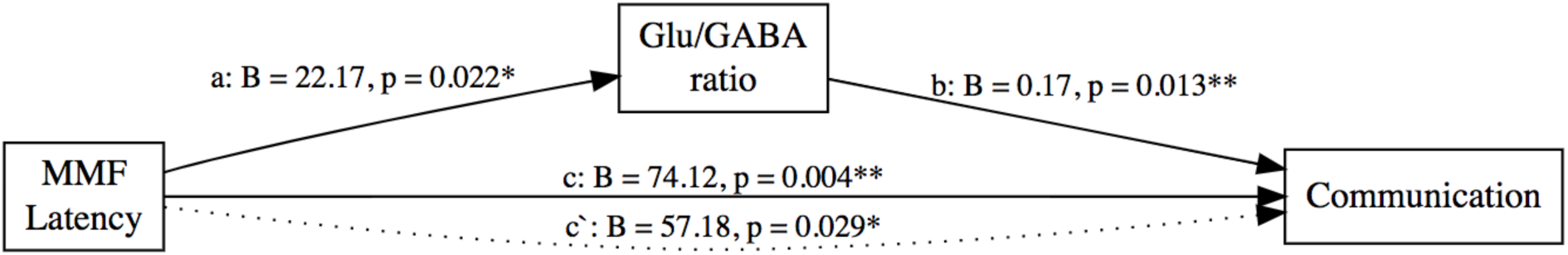
The mediation model for AQ communication. The mediating effect of glutamate (Glu)/GABA+ ratio on the relationships between mismatch field (MMF) latency and communication subscale scores from the autism spectrum quotient (AQ). The average causal mediation effect (AMCE) was not statistically significant (ß = 16.94, *p* = .161). *a =* unstandardised regression coefficient between predictor and mediator, *b* = unstandardised regression coefficient between mediator and dependant variable, *c* = unstandardised regression coefficient between predictor and dependant variable (total effect), and *c‘* = unstandardised regression coefficient between predictor and dependant variable when controlling for mediators (direct effect). *\* p* < .05, ** *p* < .01, *** *p* < .001.

Figure 4 illustrates the mediation model for SPQ interpersonal features. When controlling for age, sex and RAPM scores, MMF latency significantly predicted interpersonal features (ß = 153.85, *t* = 2.07[74.15], *p* = .047), and glutamate/GABA+ ratio (ß = 22.17, *t* = 2.43[9.14], *p* = .022). When controlling for the effect of glutamate/GABA+ ratio on this marginal relationship, the relationship between MMF latency and interpersonal features Beta coefficient reduces substantially, and is no longer significant, suggesting that glutamate/GABA+ ratio fully mediates the relationship between MMF latency and Interpersonal features (ß = 78.0, *t* = 1.40[75.31], *p* = .310). Nonparametric causal mediation analysis indicated that this mediating effect (AMCE) approached significance (ß = 75.86, 95%CI[-1.21, 170.80], *p* = .054).

**Figure 4.**
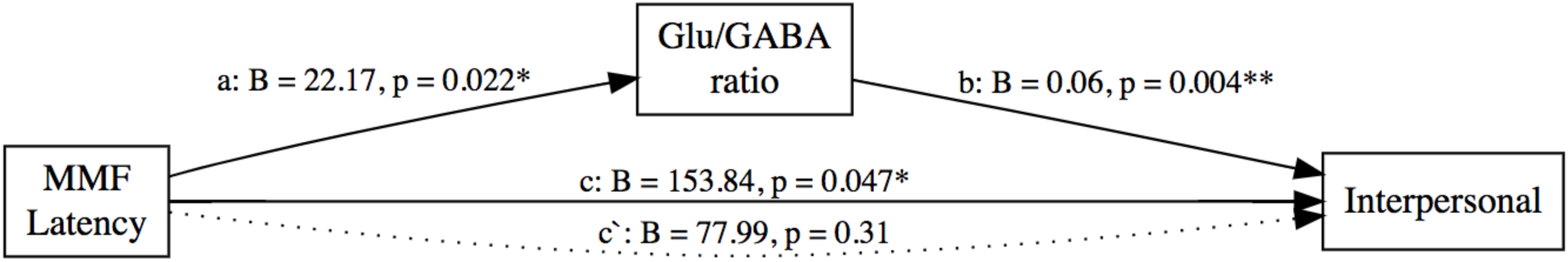
The mediation model for SPQ interpersonal features. The mediating effect of glutamate (Glu)/GABA+ ratio on the relationships between mismatch field (MMF) latency and interpersonal features from the schizotypal personality questionnaire (SPQ). The average causal mediation effect (AMCE) was approaching statistically significant (ß = 75.86, *p* = .054). *a =* unstandardised regression coefficient between predictor and mediator, *b* = unstandardised regression coefficient between mediator and dependant variable, *c* = unstandardised regression coefficient between predictor and dependant variable (total effect), and *c‘* = unstandardised regression coefficient between predictor and dependant variable when controlling for mediators (direct effect). *\* p* < .05, ** *p* < .01, *** *p* < .001.

## Discussion

This study is the first to investigate the effect of glutamate/GABA+ ratios in auditory processing regions on the relationship between MMF and symptom domains of the autism and schizophrenia spectra. Initial correlational analyses provide preliminary evidence to suggest an interrelationship between right STC glutamate/GABA+ ratios, right MMF latency and symptom domains associated with psychosocial function, such that longer MMF latency was associated with more severe psychosocial deficits, and higher glutamate/GABA+ ratios. Subsequent mediation models suggested that right hemisphere glutamate/GABA+ ratios played a mediating role in the relationship between longer MMF latency and psychosocial dysfunctions. These findings support the vast majority of existing literature pointing toward hyper-glutamatergic and/or hypo-GABAergic processes across the autism and schizophrenia spectra (Cellot and Cherubini, 2014; Deidda et al., 2014; Fatemi, 2008; Fatemi et al., 2009; Foss-Feig et al., 2017; Rojas, 2014), and suggests this might be specific to psychosocial dysfunction.

Glutamate/GABA+ ratios were shown to fully mediate the relationship between right MMF latency and schizotypal interpersonal features, while only partially mediating the relationship with autistic communication deficits. This difference in effect could be attributed to communication deficits being a more specific symptom domain, which is more closely related to the development of speech and the ability to use and understand spoken language, as well as non-verbal cues (American Psychological Association, 2013). Schizotypal interpersonal features on the other hand encompass a broader range of interpersonal domains, such as excessive social anxiety, reduced affect, and few close relationships. Interpersonal features map closely to the enduring negative symptoms characteristic of schizophrenia, which include diminished emotional expression and reduced motivation for self-initiated activities (American Psychological Association, 2013). These findings suggest that aberrant excitatory-inhibitory cycling is more closely associated with a broad range of psychosocial difficulties, rather than communication difficulties specifically. This finding is in line with several studies demonstrating aberrant excitatory-inhibitory processes associated with psychosocial difficulties in schizophrenia and autism (Ajram et al., 2019; Canitano and Pallagrosi, 2017; Foss-Feig et al., 2017; Greenwood et al., 2018; Yizhar et al., 2011).

Although the glutamate/GABA+ ratios were shown to fully mediate the relationship between MMF latency and schizotypal interpersonal features, and partially mediated the relationship with autistic communication deficits, there was no such effect for autistic social skills. This was due to the absence of a relationship between social skills and MMF latency. Somewhat surprisingly, significant relationships were observed between all AQ subscales and right MMF latency except for social skills, while only communication and social skills deficits were associated with increased glutamate/GABA+ ratios. Furthermore, moderate-strong relationships were observed between MMF latency and all three schizotypal dimensions: cognitive-perceptual features, Interpersonal features, and disorganised features, while only interpersonal features were associated with increased glutamate/GABA+ ratios. These findings suggests that increased glutamate/GABA+ ratios might be more specific to psychosocial difficulties (Foss-Feig et al., 2017), and that the neurobiological processes involved in MMF generation may contribute more broadly to autism and schizophrenia symptom domains.

The right hemisphere effects observed herein can be interpreted in relation to findings that the right hemisphere is particularly involved in social, emotional, and language processing (de Achával et al., 2012; Lindell, 2006). Furthermore, MMN/F and glutamate/GABA+ ratios have previously been associated with psychosocial deficits (Cochran et al., 2015; Ford et al., 2017a; Greenwood et al., 2018; Han et al., 2014; Hegarty et al., 2018; Yizhar et al., 2011). The effect observed here is also consistent with previous research reporting right hemisphere effects during social cognition and emotion tasks (Abu-Akel et al., 2017; de Achával et al., 2012).

The case for disrupted excitatory-inhibitory cycling across autism and schizophrenia spectrum disorders has gained significant momentum in recent years, with several lines of evidence from genetic, neuroscientific, and pharmacological studies converging to indicate multiple contributors to increased excitation-inhibition. A recent review by Ajram and colleagues (2019) detailed such converging evidence in autism, highlighting many genetic contributions to glutamate and GABA signalling pathways, such as mutations on neuroligin 3 and neurexin 1, polysynaptic density protein 95, SHANK3, CNTNAP2, and GABA_A_ receptor subtype genes. In a similar vein, there are several means by which alterations in glutamine-glutamate-GABA metabolism could lead to an excitation-inhibition imbalance, with several of the enzymes involved in glutamine-glutamate-GABA metabolism reported to be compromised in autism (e.g., glutamic acid decarboxylase 65 and 67, glutaminase; Ajram et al., 2019; Fatemi et al., 2002; Shimmura et al., 2013). Highly variable methodologies and participant sample characteristics in human 1H-MRS studies also contribute to the inconsistencies reported in regional glutamate and GABA concentrations (for review see Ajram et al., 2019; Ford and Crewther, 2016), although, these inconsistencies could be attributed to actual regional differences in excitation-inhibition (Foss-Feig et al., 2017).

Research has also reported various genetic mutations that may affect excitatory-inhibitory processes in schizophrenia, including those seen in autism (e.g., CNTNAP2, SHANK3, neurexin 1), as well as disruption to the glutamate-GABA metabolism enzymes glutaminase and glutamine synthetase (Bruneau et al., 2005). 1H-MRS studies have relatively consistently shown increased glutamate, glutamine and Glx across brain regions, which have been associated with negative symptoms (for a review, see Foss-Feig et al., 2017). Neuroscientific evidence from a mouse study demonstrated that ontogenetically increased prefrontal excitation/inhibition increased gamma band power and impair social functioning (Yizhar et al., 2011), while benzodiazepines, which modulate GABA_A_ receptors to increase inhibition, have been shown to improve social deficits in mice (Han et al., 2014). Human studies involving transmagnetic stimulation (TMS) have also demonstrated GABA_A_ receptor deficits in ASD (Enticott et al., 2013, 2010), and GABAergic modulation to improve movement-related symptoms (Enticott et al., 2012) social outcomes in mice (Tan et al., 2018). Furthermore, reduced GABA and increased glutamate concentrations have been associated with symptom severity, and increased glutamate/GABA+ ratios have been associated with poor social functioning in healthy controls (Ford et al., 2017a), thus future work would benefit from focussing on symptom specific aberrances.

Of note was the absence of a relationship between MMF amplitude and glutamate/GABA+ ratios. Previous clinical studies have reported mixed results when investigating MMN amplitude and glutamate and/or GABA concentrations in psychiatric (Chitty et al., 2015; Kaur et al., 2019; Rowland et al., 2016) and healthy populations (Kompus et al., 2015). One study reported increased Glx concentrations in the superior temporal region were associated with slightly smaller MMN amplitude in healthy populations, however the MMN was recorded fronto-centrally (Kompus et al., 2015). This study is the first to report the correlation between MEG-measured MMF and glutamate/GABA+ ratios, and thus may identify a more reliable and localised effect.

There was also no relationship observed between left or right MMF amplitude and severity of schizophrenia spectrum domains, which is somewhat surprising given EEG consistently demonstrates reduced MMN in schizophrenia samples; however, the exact mechanisms of this reduction remains unknown (Kargel et al., 2014; Michie et al., 2016; Todd et al., 2008, 2003). The MMN deficit in schizophrenia has been explained in the context of a predictive coding deficiency, whereby the reduced MMN amplitude represents a neurobiological inefficiency in forming prediction about the environment to reduce prediction error (Friston, 2005; Michie et al., 2016). A more recent meta-analysis has questioned this theory after reporting no relationship between positive and negative symptoms of schizophrenia and MMN amplitude, suggesting that although MMN is consistently reduced in schizophrenia and at-risk groups, this is independent of symptom severity (Erickson et al., 2017); MMN latency was not investigated. Post-mortem studies have reported reduction NR1 mRNAs and proteins in schizophrenia, which are essential for NMDAr functionality, and may thus explain the reduced MMN in schizophrenia samples, but this might not translate to a relationship between symptom severity and MMN amplitude (Michie et al., 2016). The findings herein suggest that the MMN elicited from the auditory cortex at the initial processing stage may not be affected by schizotypal or autism trait severity.

It should be noted that the sample size of this study was quite small, with a number of participants’ data rejected due to poor quality; thus, the effects observed should be interpreted as a preliminary finding that requires further exploration. In addition, self-report measures, such as the AQ and SPQ, require a degree of introspection that is often missing for those with poor psychosocial functioning. The AQ and SPQ tend to ask participants to report on their personal preferences, however, rather than to rate their own abilities (Baron-Cohen et al., 2001). For this study, we recorded SPQ data on a 4-point Likert scale as per AQ, and pseudo-randomised items in an attempt to reduce response bias.

Clearly, the hypothesis of abnormal excitation-inhibition cycling in autism and schizophrenia is far from straightforward, and further multimodal research is necessary to better understand the complexity of the pathways contributing to the excitation-inhibition in psychopathology (Foss-Feig et al., 2017). This study utilised two neurological models of excitatory-inhibitory processes using two neuroimaging modalities: MMF with MEG, and glutamate and GABA concentrations with 1H-MRS. The combined use of MEG and 1H-MRS has allowed for the relationship between electrophysiological processes and behavioural outcomes to be better understood in relation to the effect of regional glutamate and GABA concentrations – a direct measure of excitatory and inhibitory processes.

## Acknowledgements

We thank Swinburne Neuroimaging for their support through data collection and analysis. The authors also acknowledge the facilities of Swinburne Neuroimaging (SNI) and its flagship funding from the Australian National Imaging Facility (NIF) under the National Collaborative Researcher Infrastructure Strategy (NCRIS) implemented by the Australian Government.

