## Supplementary for "Cortical excitation-inhibition ratio mediates the effect of pre-attentive auditory processing deficits on interpersonal difficulties"

**Supplementary Information**

Supplementary Table 1. 1H-MRS fit statistics for left and right superior temporal voxel.

| | Left | | Right | |
|---|---|---|---|---|
| | Mean(*SE*) | Range | Mean(*SE*) | Range |
| Grey Matter | 66.09 (1.0) | 50.62 - 73.54 | 66.84 (0.94) | 49.19 - 74.36 |
| White Matter | 26.04 (1.26) | 16.03 - 47.86 | 24.06 (1.23) | 13.98 - 46.02 |
| Cerebrospinal Fluid | 7.87 (.48) | 1.52 - 15.46 | 9.1 (0.48) | 4.15 - 15.37 |
| Glu water FWHM (Hz) | 7.58 (.19) | 6.05 - 10.54 | 8.35 (0.25) | 5.78 - 11.52 |
| Glu water Frequency SD (Hz) | .35 (.24) | -2.05 - 2.78 | -0.26 (0.44) | -11.86 - 3.52 |
| Glu SNR | 46.68 (1.74) | 30.08 - 67.12 | 48.58 (1.92) | 27.37 - 81.07 |
| Glu CRLB | 1.22 (.09) | .50 - 2.86 | 0.96 (0.1) | 0.41 - 2.73 |
| Gln CRLB | .58 (.04) | .21 - 1.46 | 0.48 (0.05) | 0.18 - 1.39 |
| Glx water FWHM (Hz) | | | | |
| Glx water Frequency SD (Hz) | | | | |
| Glx SNR | | | | |
| Glx CRLB | | | | |
| GABA Creatine FWHM (Hz) | 9.06 (.16) | 7.77 - 11.87 | 9.03 (0.14) | 7.2 - 11.19 |
| GABA water Frequency SD (Hz) | .80 (.12) | .21 - 3.98 | 1.04 (0.19) | 0.19 - 5.75 |
| GABA Fit Error | 10.78 (.48) | 4.54 - 15.81 | 8.7 (0.41) | 5.37 - 15.46 |
| GABA SNR | 9.98 (.55) | 6.33 - 22.02 | 12.3 (0.56) | 6.47 - 18.63 |

Notes: SE = standard error, Glu = glutamate, Gln = glutamine, Glx = glutamate + glutamine, FWHM = full-width half maximum, SD = standard deviation, SNR = signal to noise ratio, CRLB = Cramer-Rao lower bound.

Supplementary Table 2. Spearman's rank order (ρ) correlations between glutamate, Glx and GABA, and MMF amplitude and latency for left and right hemisphere.

| | N = 33 | Glutamate | Glx | GABA+ | MMF amplitude |
|---|---|---|---|---|---|
| Left | | | | | |
| | Glx | .88** | | | |
| | GABA+ | -.03 | .02 | | |
| | MMF amplitude | .28 | .18 | .33 | |
| | MMF latency | .04 | .08 | .14 | -.12 |
| Right | | | | | |
| | Glx | .90** | | | |
| | GABA+ | -.07 | -.12 | | |
| | MMF amplitude | -.03 | .15 | -.15 | |
| | MMF latency | -.15 | -.10 | -.38* | .15 |

Notes: *$p < .05$, **$p < .001$. Glx = glutamate + glutamine, GABA+ = GABA + macromolecules, MMF = mismatch field

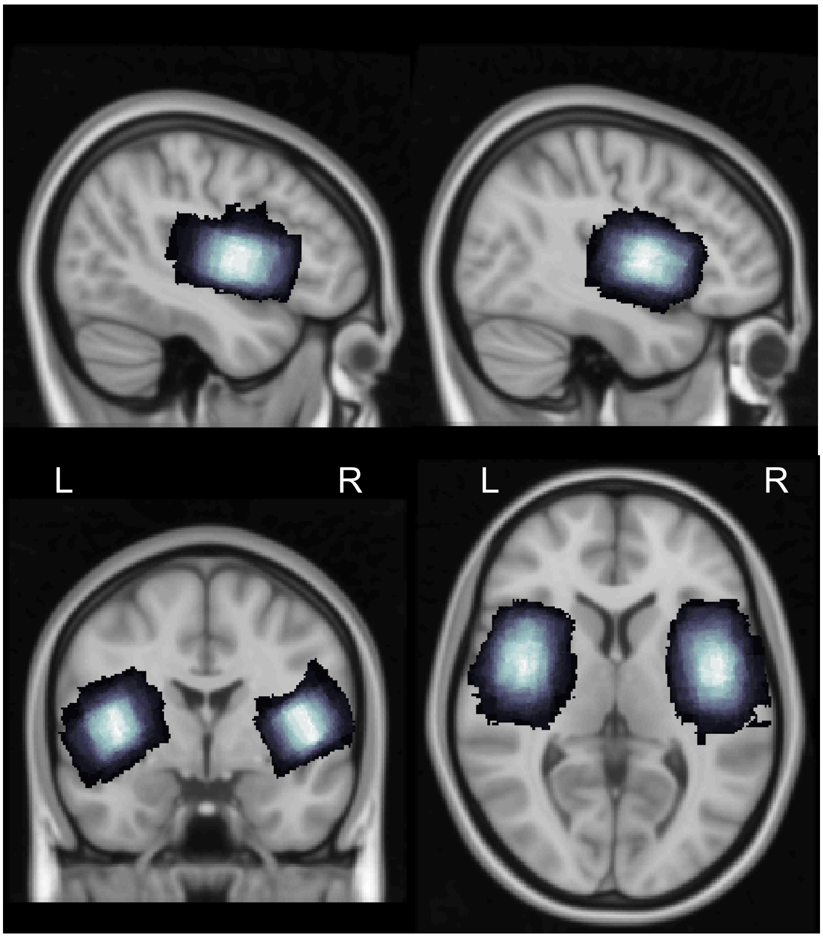

Supplementary Figure 1: Mean voxel placement of left and right superior temporal MRS voxel on standard MNI structural image. Left and right voxels in the sagittal slice are shown at top left and top right, respectively. Lighter colours represent more overlapping voxels.
